# Reliable method for high quality His-tagged and untagged *E. coli* phosphoribosyl phosphate synthase (Prs) purification

**DOI:** 10.1101/864173

**Authors:** Walter Beata Maria, Szulc Aneta, Glinkowska Monika

**Affiliations:** Department of Bacterial Molecular Genetics, University of Gdansk, Wita Stwosza 59, Gdansk, Poland

**Keywords:** *Escherichia coli*, Phosphoribosyl phosphate synthase, Prs, PRPP, activity, Temperature-sensitive *prs-2* mutant

## Abstract

Prs (phosphoribosyl pyrophosphate synthase) is a broadly conserved protein that synthesises 5-phosphoribosyl 1-pyrophospate (PRPP); a substrate for biosynthesis of at least 10 enzymatic pathways including biosynthesis of DNA building blocks – purines and pyrimidines. In *Escherichia coli*, it is a protein of homo-hexameric quaternary structure, which can be challenging to work with, due to frequent aggregation and activity loss. Several studies showed brief purification protocols for various bacterial PRPP synthases, in most cases involving ammonium sulfate precipitation.

Here, we provide a protocol for expression of *E. coli* Prs protein in Rosetta (DE3) and BL21 (DE3) pLysE strains and a detailed method for His-Prs and untagged Prs purification on nickel affinity chromatography columns. This protocol allows purification of proteins with high yield, purity and activity. We report here N-terminally His-tagged protein fusions, stable and active, providing that the temperature around 20 °C is maintained at all stages, including centrifugation. Moreover, we successfully applied this method to purify two enzyme variants with K194A and G9S alterations. The K194A mutation in conserved lysine residue results in protein variant unable to synthetize PRPP, while the G9S alteration originates from *prs-2* allele variant which was previously related to thermo-sensitive growth. His-PrsG9S protein purified here, exhibited comparable activity as previously observed *in-vivo* suggesting the proteins purified with our protocol resemble their physiological state.

The protocol for Prs purification showed here indicates guidance to improve stability and quality of the protein and to ensure more reliable results in further assays *in-vitro*.

## INTRODUCTION

Prs catalyzes formation of PRPP by transfer of β,γ-pyrophosphate from ATP to C1 hydroxyl group of ribose 5-phosphate (R5P), yielding AMP as a second product [1]. PRPP is a crucial compound in biosynthesis pathways of purine and pyrimidine nucleosides, pyridine enzyme cofactors like nicotinamide adenine dinucleotide (NAD), as well as amino acids histidine and tryptophan [2, 3]. Phosphoribosyl pyrophosphate synthases are ubiquitous and conserved enzymes, with overall identity of amino acid sequence between the respective *E. coli* and human enzymes reaching 47% [4]. Three classes of PRPP synthases are distinguished, based on their quaternary structure, inorganic phosphate requirement, ability to utilize nucleotide triphosphates other than ATP as pyrophosphate donors and presence of allosteric regulation site [5-9]. *E. coli* Prs protein, as well as their homologs of many other bacteria and mammals, belongs to Class I of PRPP synthases. This class is characterized by hexameric quaternary structure, whereas their activity is allosterically regulated by ADP and GDP (guanosine diphosphate). These PRPP synthases utilize ATP exclusively as a diphosphate donor [9-12]. Crystal structure of *E. coli* PRPP synthase has been resolved revealing propeller-shaped trimmer of dimers [12, 13]. C-terminal domains of Prs form the blades of the propeller, whereas the core is comprised of N-terminal domains [12, 13]. ATP-binding site in the catalytic centers of Prs is composed of residues of two chains (F35, D37, E39 of one chain and R96, Q97, D98, R99 and H131 of the second chain). Ribose 5-phosphate on the other hand is coordinated by C-terminal residues of a single chain (D224, T225, G226, K231) [13]. Allosteric regulatory site of *E. coli* PRPP synthase is distal from the catalytic center and made up by residues of three chains at the propeller core (chain 1: Q136, D144, S308, S310, F313; chain 2: R49, S81; chain 3: R102, S103, R105). Allosteric inhibition by ADP is competitive with stimulation of Prs activity by phosphate and magnesium ions, suggesting that they all bind at the same site [9, 11, 14, 15]. Moreover, several residues form additional intra-molecular interactions between hexamer chains, among them is conserved Lys194, essential for the activity of *E. coli* PRPP synthase [16, 17].

Several viable *E. coli prs* mutants have been isolated, among them *prs-2* which displays a temperature-sensitive phenotype and is unable to grow at temperatures above 25°C [2]. The *prs-2* allele carries two mutations, one resulting in G9S alteration of the protein’s primary structure and another one located in the non-coding region near the Shine-Dalgarno sequence. The former change in conserved glycine residue reduces activity of the enzyme and is responsible for its heat-liability. The latter – results in increased enzyme synthesis in *E. coli* cell [18]. The exact role of Gly9 residue is unknown.

Although PRPP synthases are well-characterized biochemically, regulation of their activity *in-vivo* remains poorly understood. For instance, *K*_*m*_ values for bacterial Prs proteins are an order of magnitude lower than intracellular PRPP concentration, meaning that PRPP is not the rate-determining factor for those enzymes [19]. In bacteria, PRPP level is reduced in the presence of exogenous purines [20-22] and ADP-dependent allosteric inhibition of the synthase is most likely responsible for this effect. Further studies on physiological regulation of PRPP pools and Prs activity will nevertheless require isolation of the enzyme and its variants and performance of biochemical assays estimating its activity. Prs is a complex protein, challenging to work with, as it easily aggregates and loses activity upon isolation. Several reports presented purification protocols for various bacterial PRPP synthases [6, 9, 12], however many of them involve salting out of the protein with ammonium sulfate or precipitation. In this report we provide a detailed protocol for isolation of *E. coli* Prs protein with high yield, purity and activity. Using the same protocol, we also purified two variants of the enzyme, bearing K194A and G9S alterations.

## MATERIALS AND METHODS

Media, antibiotics and buffer components used in this study were purchased from either Carl Roth or BioShop Life Science. Primers were synthesised by SigmaAldrich/Merck. Polymerases and enzymes used for cloning were purchased from Thermo Scientific or New England Biolabs.

### Bacterial strains and plasmids

*E. coli* DH5α was used for recombinant plasmid construction. *E coli* Rosetta (DE3) and *E. coli* BL21 (DE3) pLysE were used as protein expression hosts. The plasmid pET28a (EMD Biosciences) and our modified plasmid pET28a-TEV, were used as expression vectors (Table 1). For pET28a-TEV, we replaced T7 tag and partial MCS with TEV cleavage site to produce a universal expression vector allowing N-terminal (Figure 1B) or C-terminal (Figure 1C) cleavable His_6_-tagging.

**Table 1.** Plasmids in this study.

| Plasmid | Features | Source |
| --- | --- | --- |
| pET28a | KanR, T7 promoter, 6xHis, T7-tag, MCS | EMD Biosciences |
| pET28a-TEV | KanR, T7 promoter, 6xHis-TEV-6xHis | This study |
| pET28a-His-Prs | KanR, T7 promoter, 6xHis tag on N-terminus of Prs | This study |
| pET28a-His-TEV-Prs | KanR, T7 promoter, 6xHis tag cleavable with TEV protease on N-terminus of Prs | This study |
| pET28a-His-PrsK194A | KanR, T7 promoter, 6xHis tag on N-terminus of Prs with point mutation K194A | This study |
| pET28a-His-PrsG9S | KanR, T7 promoter, 6xHis tag on N-terminus of Prs with point mutation G9S | This study |

**Figure 1.**
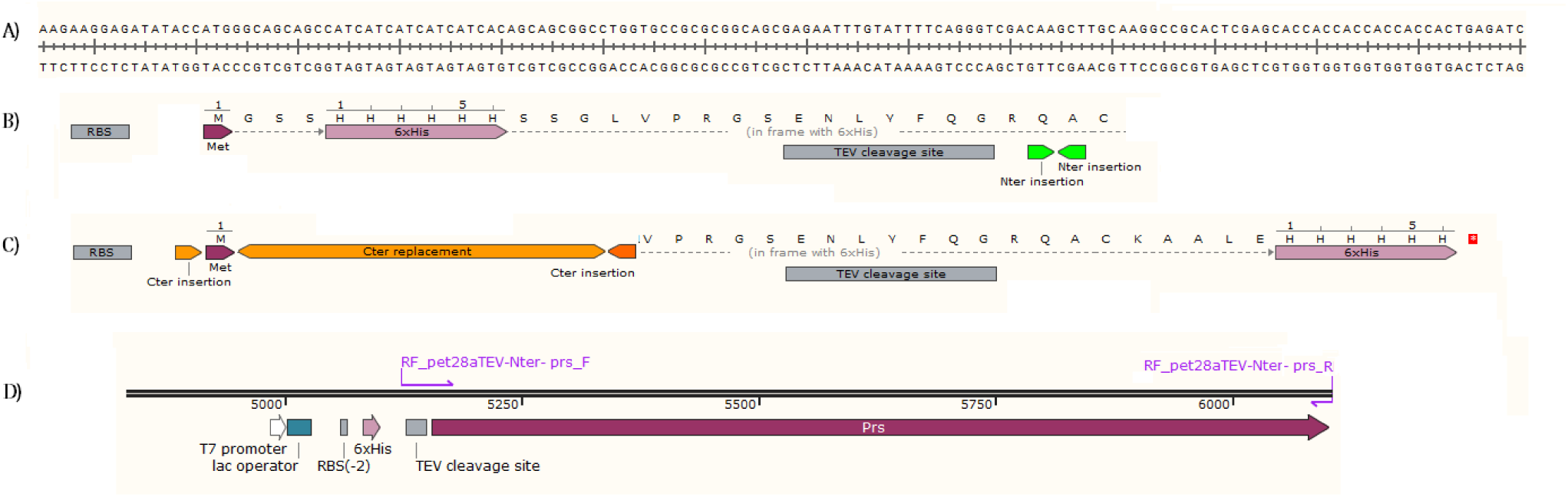
Partial map of the pET28a-TEV plasmid. A) DNA sequence of pET28a-TEV cloning site, B) RF cloning strategy for N-terminally His_6_-tagged proteins, C) RF cloning strategy for C-terminally His_6_-tagged proteins, D) Fragment of pET28a-His-TEV-Prs plasmid.

### Molecular cloning and construction of recombinant plasmids

Two pET28a plasmids carrying N-terminally His_6_-tagged *prs* and N-terminally His_6_-tagged *prs* with TEV cleavage site were cloned (Table 1) with restriction free cloning method [23]. Primer pair RF_pET28a-prs_F and RF_ pET28a-prs_R was used for for pET28a-His-Prs vector and primer pair RF_pET28a-TEV-prs_F and RF_pET28a-TEV-prs _R for pET28a-His-TEV-Prs vector (Table 2). Briefly, the mega primers were amplified on *E. coli* MG1655 genomic DNA with Phusion Flash High-Fidelity PCR Master Mix (Thermo Scientific), the PCR product was purified with NucleoSpin Gel and PCR clean-up kit according to manufacturer’s description (Macherey-Nagel, 740609). RF cloning into pET28a was performed with eighteen 3-step cycles with annealing temperature 65°C and extension time 7 minutes. RF cloning into pET28a-TEV was more efficient when 2-step cycling was applied. Following thermal cycling, the template plasmid was digested with DpnI at 37°C overnight, and remaining DNA was used for transformation of chemically competent DH5α and plated on LB supplemented with 50 µg/ml kanamycin. The colonies were screened with colony PCR using primers pair s-pET28a_F and s-pET28a_R (Table 2) and confirmed with sequencing.

**Table 2.**
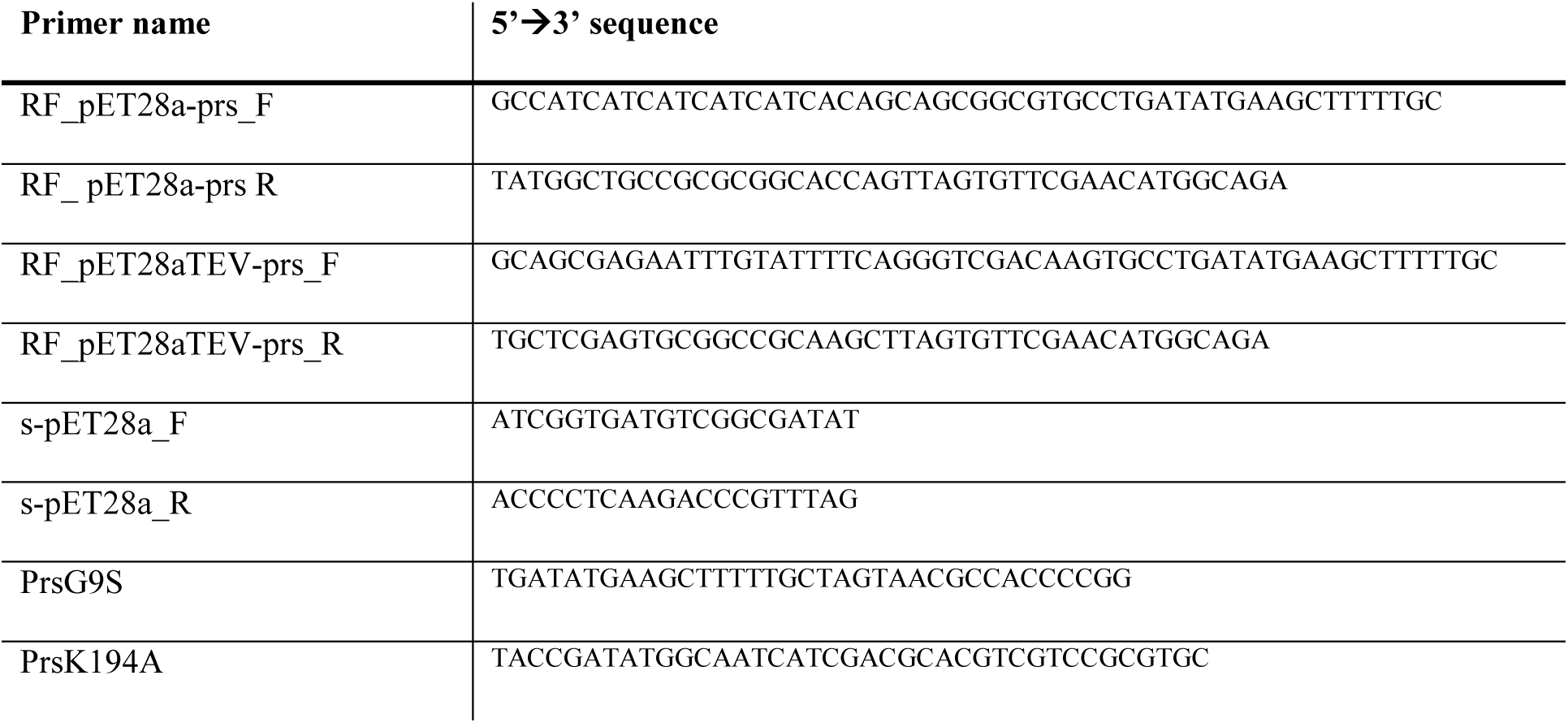
Primers used in this study.

In addition, we cloned two pET28a-His-Prs plasmid variants (Table 1). One of them overproducing Prs with single amino acid substitution replacing conserved lysine K194 with alanine (plasmid pET28a-His-PrsK194A). Another plasmid carried thermo-sensitive *prs* allele resulting in replacement of glycine G9 residue with serine (*prs-2* mutant, plasmid pET28a-His-PrsG9S). Point mutations were introduced into the vector sequences with phosphorylated primers K194A and G9S (Table 2). Phosphorylation of primers (10 µM) was done with ATP (1mM), T4 Polynucleotide Kinase (10 U), 1x PNK buffer A in 50 µl reaction, incubated at 37°C, for 30 min, followed by 20 min inactivation at 65°C. Phosphorylated primer (0.48 µM) with 12.5 µl Phusion Flash Master Mix, Taq ligase (20 U), 1 × Taq ligase buffer, and 60 ng of pET28a-His-Prs in a 25 µl reaction, was used in 30 PCR cycles with annealing at 55°C and 7 minutes extension at 65°C. The PCR reaction was treated with DpnI and used for transformation of chemocompetent DH5α as described above.

### Overexpression of *prs* in *E. coli* strains

Chemocompetent *E coli* Rosetta (DE3) were transformed with plasmids pET28a-His-Prs, pET28a-His-PrsK194A and pET28a-His-TEV-prs while *E. coli* BL21 (DE3) pLysE was transformed with pET28a-His-PrsG9S. Overnight culture in LB supplemented with 50 µg/µl kanamycin and 34 µg/µl chloramphenicol were diluted 1:100 into 1 L of 2xYT medium (BioShop Life Science) supplemented with antibiotics and grown at 37°C until OD_600_ reached 0.8-1. Bacterial Cultures were induced with 100 µM Isopropyl β-D-1-thiogalactopyranoside (IPTG) except for *E. coli* BL21 (DE3) pLysE::pET28a-His-PrsG9S which was induced with 1 mM IPTG. After induction, cultures were incubated at 37°C for another 5 h, pelleted by centrifugation and frozen at −20°C.

### Purification of His-Prs, His-PrsK194A and His-PrsG9S

**Buffer A**: 50 mM potassium phosphate buffer (pH 7.5), 10 % glycerol, 500 mM NaCl, 20 mM imidazole (pH 7.8);

**Lysis buffer:** buffer A supplemented with 0.1% Tween 20. A pinch (1-2 mg) of lysozyme from chicken egg white was added after the pellet was suspended in the buffer prior sonication;

**Buffer B**; 50 mM potassium phosphate buffer (pH 7.5), 10 % glycerol, 500 mM NaCl, 300 mM imidazole (pH 7.8), 0.5 mM tris(2-carboxyethyl)phosphine (TCEP);

**SEC1 buffer**; 50 mM potassium phosphate – NaOH buffer (pH 8.2), 10 % glycerol, 500 mM NaCl;

**SEC2 buffer**; 50 mM potassium phosphate buffer (pH 7.5), 10 % glycerol, 200 mM NaCl, 1mM MgCl_2_;

**SEC3 buffer**; 50 mM potassium phosphate buffer (pH 7.5), 10 % glycerol, 150 mM L-Glutamic acid monopotassium salt.

Cells were disrupted in lysis buffer on ice by 7 cycles of 1 minute sonication with pulsing, with 1 minute interval to avoid overheating. Lysates were cleared by centrifugation for 1 h at 32 000 × g at 10 °C. Proteins were purified by Ni-affinity chromatography on 5 mL His-Trap columns (GE Healthcare) as follows: the His-Trap columns were equilibrated with 10 volumes of buffer A and the cleared lysates were loaded on a column followed by >10 volumes buffer A wash. Further, the columns were washed with 15 mL 5 % buffer B in buffer A followed by wash with 20 mL 10 % buffer B in buffer A and 20 mL 15 % buffer B in buffer A. Proteins were eluted in buffer B as 6 fractions, 5 mL each. Elution fraction number 2 containing highest protein concentration was immediately diluted with 15 mL of SEC1 to decrease imidazole concentration below 100 mM. Elution fractions number 3 and 4 were diluted with 10 mL of SEC1 buffer while elution fractions number 5 were diluted with 5 mL of SEC1 buffer to decrease imidazole concentration down to 100-150 mM. Proteins were handled at 18-20 °C at all times.

Protein fractions were combined and dialysed overnight at 19 °C. Three buffers were tested SEC1, SEC2 and SEC3 in three independent dialyses. Further, the proteins were concentrated with Amicon Ultra 4 MWCO10kDa (Millipore) to approximately 3 – 4 mL. Concentration was performed at 16 – 20 °C with centrifugation at 2250 × g.

Proteins were further purified by size-exclusion chromatography at 15 – 20 °C on Superdex 200 10/300 GL gel filtration column (GE Healthcare) each, in three protein loads on a column. Purified fractions were checked on a SDS-PAGE gel, concentrated as described above to approximately 10-13 mg/mL.

Protein concentrations were measured with NanoDrop Spectrophotometer (Thermo Scientific) at 280 nm. Purified proteins were snap-frozen in liquid nitrogen and stored at −70°C.

### Purification of untagged Prs

**Buffer A**; as above;

**Lysis buffer:** as above;

**Buffer B**; as above;

**Buffer A0-H pH 7.5**; 20 mM HEPES, 10 % glycerol, 2 mM MgCl_2_;

**Dialysis buffer pH 7.8**; 20 mM HEPES, 100 mM L-Glutamic acid monopotassium salt, 10 % glycerol, 1mM MgCl_2_;

S**EC1 buffer**; as above;

**SEC3 buffer**; as above.

The cell lysis and purification on His-Trap column of His-TEV-Prs protein variant was done identically as for His-Prs. However, the 5 mL eluted fractions were diluted in buffer A0-H pH 7.5. In addition, to fraction number 2, 5 mL of dialysis buffer was added to avoid precipitation. Eluted fractions were analysed on SDS-PAGE gel and selected fractions were combined and transferred into 50 mL falcon tubes as 20 mL aliquots. DTT and EDTA to a final concentration of 1 mM each were added into each aliquot. TEV protease (0.2 mg) was added to each 20 mL aliquot. The TEV protease used in this study was purified in-house from pRK793 vector in buffer 50 mM Tris-HCl pH 8.0, 200 mM NaCl, 10 % glycerol 2 mM β-mercaptoetanol. Aliquots were incubated at 24 °C for 4 h in horizontal position with no shaking or rotation. After incubation, aliquots were dialysed against dialysis buffer pH 7.8 overnight at 22 °C.

The tag and the protease were removed by Ni-affinity chromatography. Briefly, the protein was loaded on a His-Trap column, calibrated with dialysis buffer. Flow through was collected in 10 mL fractions, followed by 10 mL wash A and elution with buffer B. Flow through fractions, containing untagged Prs protein, were combined and dialysed overnight at 19 – 20 °C independently against SEC1 or SEC3 buffer.

The protein was concentrated with Amicon Ultra 4 MWCO30kDa (Millipore) at 20 °C at 1650 × g. Optionally, the protein can be further purified by size-exclusion chromatography at 15 – 20 °C on Superdex 200 10/300 GL gel filtration column (GE Healthcare) as described above.

### Measurement of Prs catalytic activity

The catalytic activity of His-Prs and untagged Prs was measured by assessing the amount of AMP produced by Prs as a second product along with PRPP formation from R5P and ATP. Briefly, 2.45 pmol of of Prs hexamer was incubated in 100 µl Prs reaction buffer (50 mM Tris, pH 8.0, 100 mM KCl, 13 mM MgCl_2_, 0.1 mg/ml BSA, 0.5mM DTT, 50 µM ribose-5-P phosphate, 50 µM ATP) for 30 minutes at 37°C with orbital shaking. His-Prs-K194A inactive variant was used as a negative control. After incubation, tubes were briefly centrifuged and 10 µl was transferred into fresh PCR tube in duplicates as a technical repeat. Remaining ATP was removed, AMP was converted to ADP and further ATP was measured through luciferase reaction with the AMP-Glo assay according to manufacturer’s description (Promega) with incubation at 24 °C. Activity of each protein was measured at least in three biological repeats.

## RESULTS AND DISCUSSION

### Purified, active His-Prs fusion protein as well as its G9S and K194A variants are stable at room temperature

His-Prs, His-PrsG9S and His-PrsK194A were successfully purified yielding up to 80 mg of pure protein, as determined by SDS-PAGE, from 1 L of medium. The novel purification protocol described here is optimised to significantly decrease protein loss due to precipitation. Substantial improvement of proteins stability in a solution was achieved by carrying out the entire purification procedure at 20 °C. Immediate dilution of proteins upon elution with buffer B minimised detrimental effect of imidazole.

Three Prs protein variants carrying non-cleavable His_6_-tag were purified on His-Trap column (Figure 2). It was observed that high concentration of imidazole is unfavourable to the proteins causing protein precipitation upon elution. This can be avoided by immediate dilution of the proteins with imidazole free buffer, in this case – buffer SEC1 used for dialysis and size exclusion chromatography. Furthermore, it was noticed, Prs protein is highly sensitive to low temperatures. Therefore, storage on ice and dialysis at 4 °C, characteristic to many protein purification procedures, results in protein loss, in its vast majority, due to precipitation. Thus, it is recommended to perform all the Prs purification procedures at temperatures around 18-20 °C, including centrifugation.

**Figure 2.**
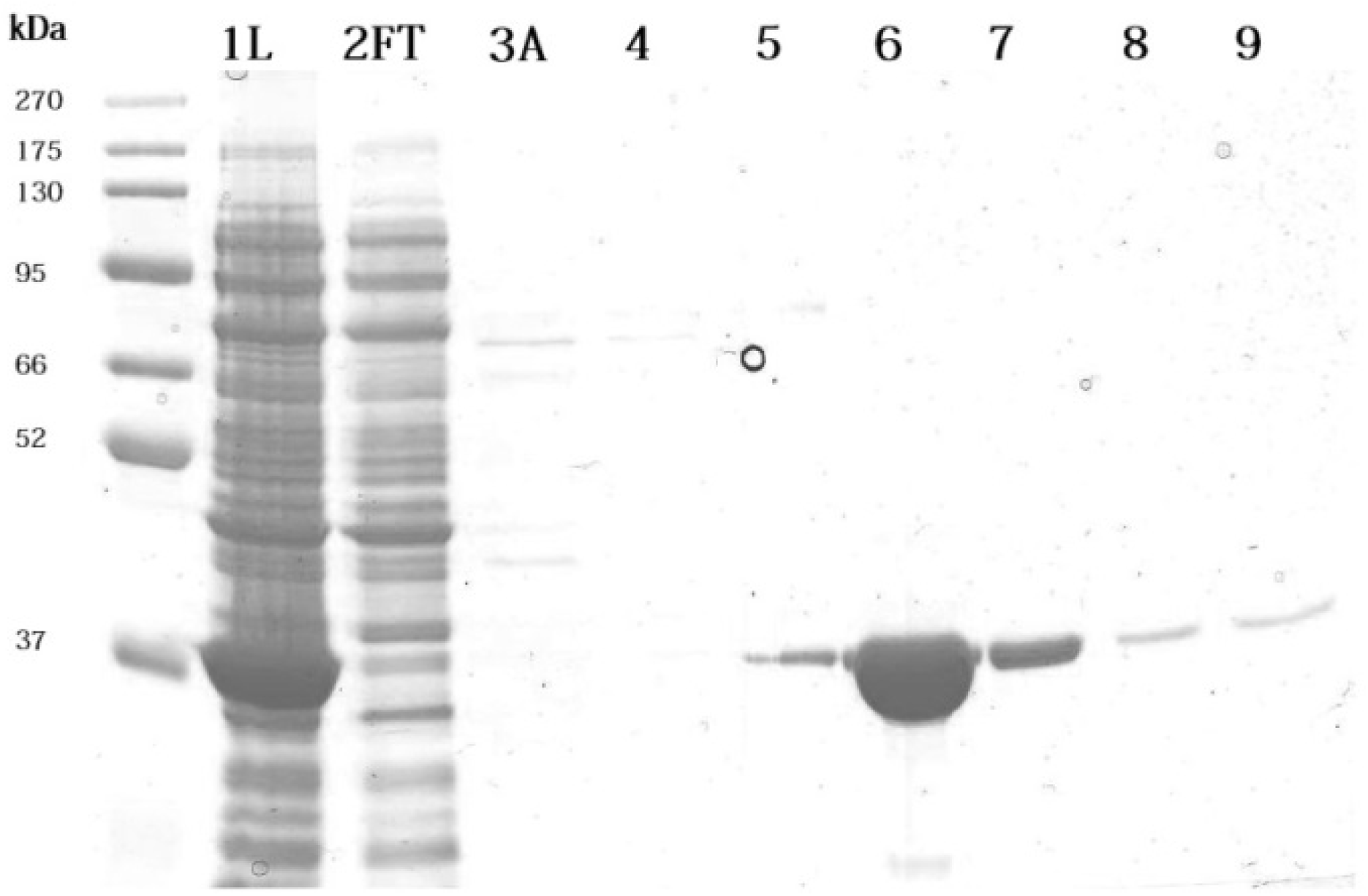
His-Prs purification on HisTrap Ni-NTA columns. His-Prs was extracted from 1 L culture of *E. coli Rosetta* (DE3) strain. **1L:** cleared lysate, **2FT:** flow through, **3A:** wash with buffer A, **4 :** final wash with 15% Buffer B in buffer A, **5 :** elution with buffer B, **6-9** : elution fractions 2, 3, 4, 5, respectively. Each fraction was diluted with 2x SDS loading dye and 10 µl was loaded on a 10 % SDS-PAGE gel. Proteins bands were visualised with Coomassie Brilliant Blue staining.

Three different buffers for final dialysis were tested; SEC1, SEC2 and SEC3 (Figure 3). His-Prs is stable at wide range (200-500 mM) of NaCl salt concentrations. Moreover, L-Glutamic acid monopotassium salt can also be used at concentration 100-150 mM providing comparable protein stability. Buffer supplementation with glycerol (10 %) is advised.

**Figure 3.**
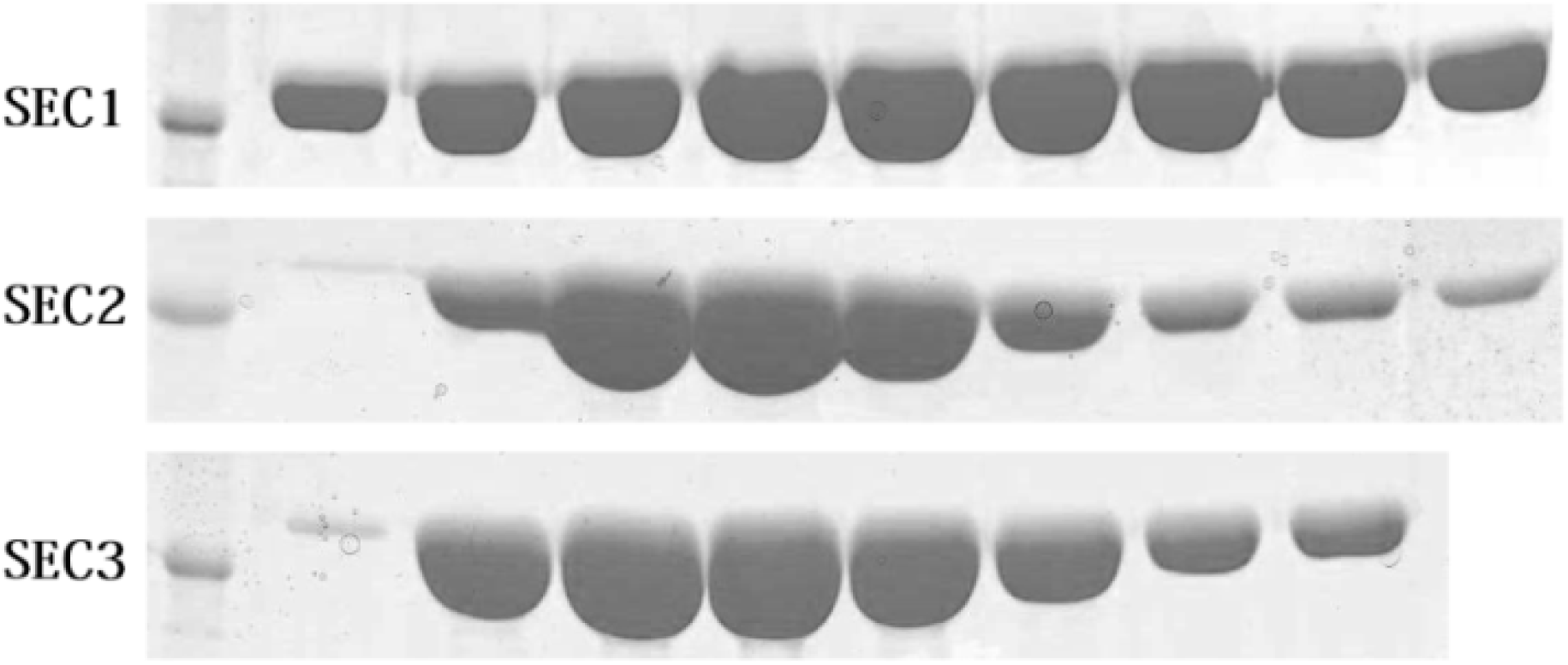
His-Prs purification – polishing step using size exclusion chromatography. Chromatography was performed on Superdex200 10/300 GL in buffers SEC1, SEC2 and SEC3. Each fraction was diluted with 2x SDS loading dye and 10 µl was loaded on a 10 % SDS-PAGE gel. After separation, proteins were visualised with Coomassie Brilliant Blue protein staining.

In addition to our observations on protein stability during protein purification, we measured if proposed storage buffers influence protein activity. His-PrsK194A protein variant carries alanine instead of lysine at position 194 resulting in loss of catalytic activity. Thus, this stable but inactive protein was used as a negative control in the activity assays. The activity of His-Prs wild type protein was measured for all three buffers proposed in this study and showed comparable results (Figure 4A).

**Figure 4.**
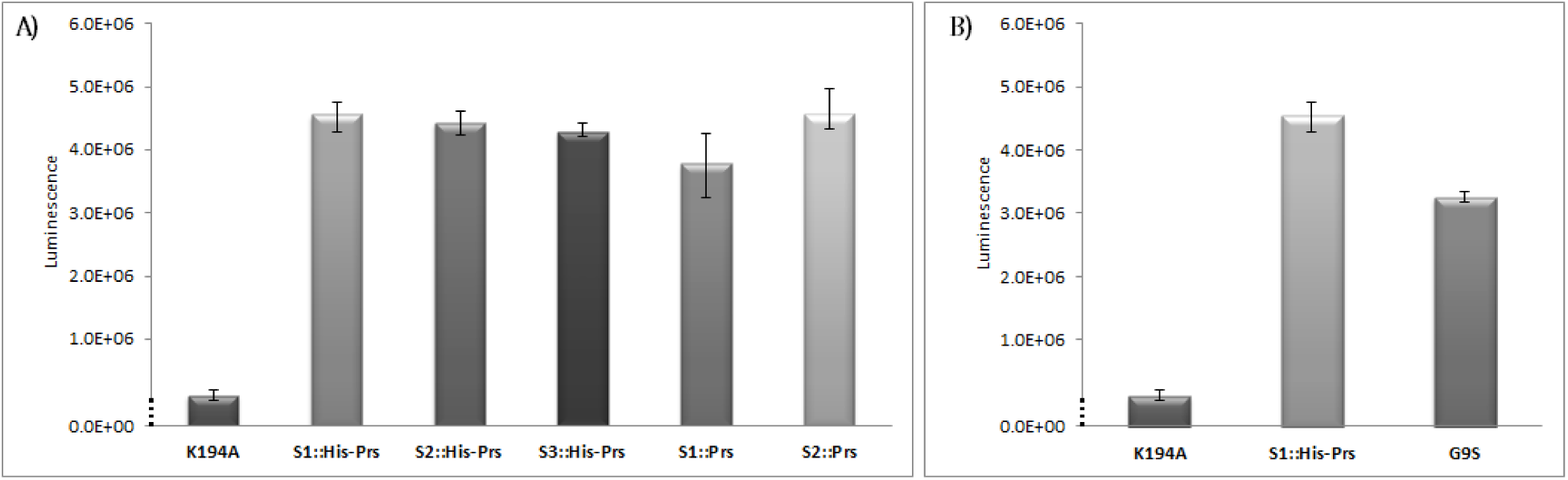
Activity of purified Prs variants. A) Comparision of His-Prs and untagged Prs activity following their purification in different buffers. K194A was purified in SEC1 buffer. S1/S2/S3 designation next to protein variant stands for SEC1, SEC2, SEC3 buffers respectively. B) Comparison of His-Prs wild type and His-PrsG9S mutant activity dialysed against buffer SEC1. The activity of Prs protein was measured in Prs reaction buffer at final hexamer concentration 24.5 nM as a luminescence signal from AMP formed from catalytic formation of PRPP from R5P (50 µM) and ATP (50 µM) in 30 minutes.

It has been shown previously that, when expressed from a plasmid and assessed *in-vivo*, PrsG9S protein variant has about half catalytic activity of the wild-type Prs [18]. Consistently, we also observed that this His-PrsG9S protein variant purified with our protocol, exhibited decreased activity in comparison to the wild-type in *in-vitro* assays (Figure 4B). Considering that His-Prs wild-type purified as described in this work exhibits high activity, we conclude that the proteins purified according to our protocol maintain their physiological characteristics what is crucial to compare the results of studies *in-vivo* with biochemical assays *in-vitro*.

### The untagged Prs is more sensitive to buffering conditions in comparison to His-Prs protein fusion

The untagged Prs was successfully purified from 1 L of *E. coli* BL21 (DE3) pLysE culture, yielding up to 50 mg of pure protein, as determined by SDS-PAGE. Untagged Prs variant also required decreasing imidazole concentration immediately after elution from Ni-affinity column, as well as maintaining 20 °C throughout the entire procedure to avoid protein aggregation.

His-TEV-Prs was successfully purified on His-Trap column yielding high amount of protein (Figure 5). The His_6_-tagged protein fusion containing TEV cleavage site was unstable in buffer SEC1 with high salt concentration (NaCl). However, it was stable at lower concentration of either NaCl or L-Glutamic acid monopotassium salt, both in HEPES and potassium phosphate buffers.

**Figure 5.**
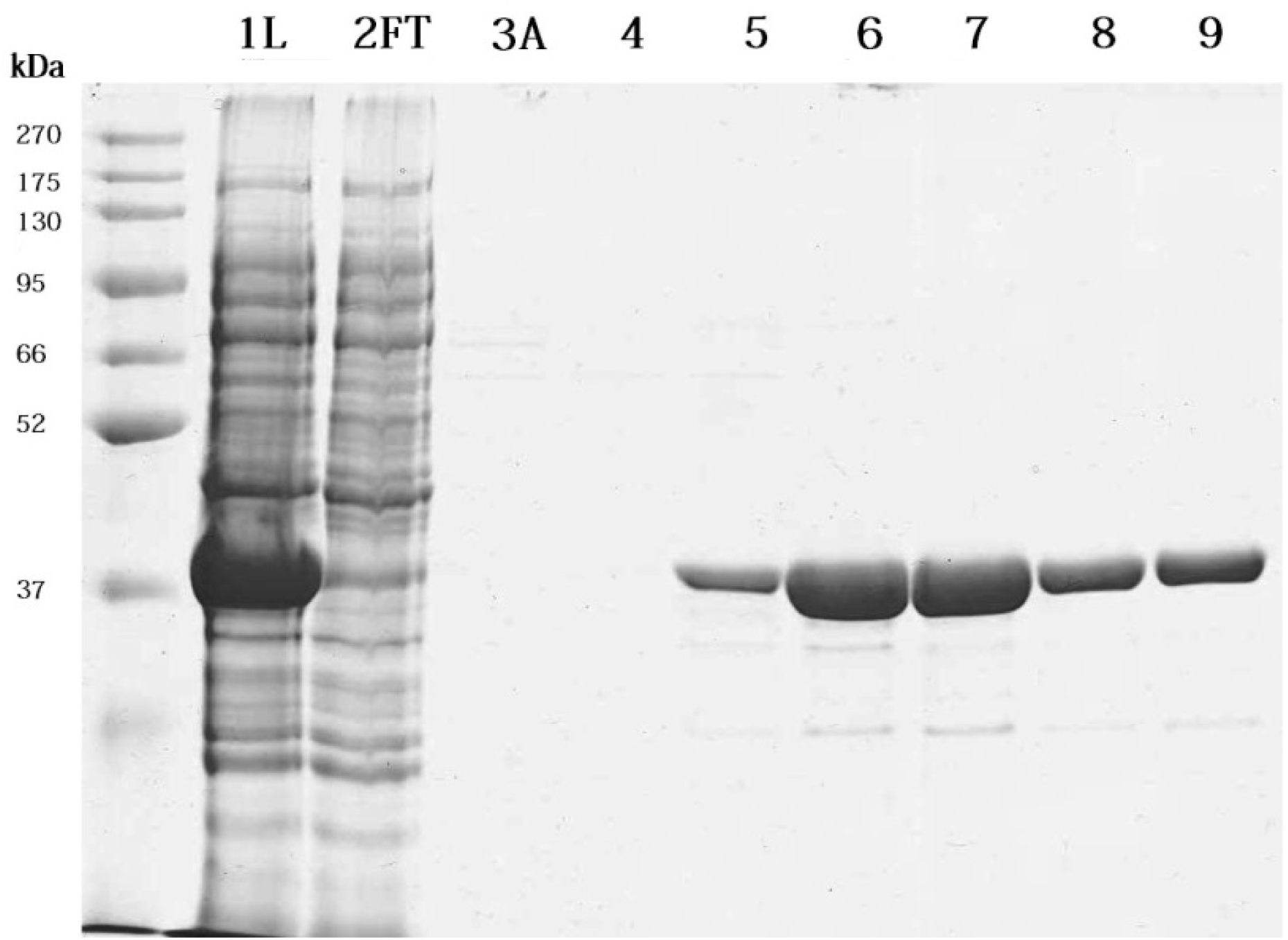
His-TEV-Prs purification using HisTrap Ni-NTA column. **1L:** cleared lysate, **2FT:** flow through, **3A**: Wash with buffer A, **4:** last wash with 15% Buffer B in buffer A, **5:** 7 mL elution B, **6:** elution-fraction 2, **7:** elution-fraction 3, **8:** elution-fraction 4, **9:** elution-fraction 5. Each fraction was diluted with 2x SDS loading dye and 10 µl was loaded on a 10 % SDS-PAGE gel. After separation, proteins on a gel were visualised with Coomassie Brilliant Blue protein staining.

Furthermore, probably due to hindered accessibility of TEV protease to N-terminally located His-TEV tag, embedded in the core of a propeller-shaped hexameric quaternary structure, cleavage with TEV protease required optimisation in order to minimise protein loss due to partial cleavage and formation of complexes consisting of both tagged and cleaved protein (Figure 6). We observed higher efficiency of TEV mediated His-TEV-Prs protein fusion cleavage in HEPES buffer, therefore it was the buffer of choice for diluting elution fractions. HEPES-based buffer did not affect Prs protein stability. Moreover, since TEV protease is more active in lower salt concentrations, we diluted the protein with HEPES based, salt free buffer A0-H. It is however important to note, dilution with A0-H buffer should be done very gently, in a drop wise manner to avoid protein precipitation. Moreover, the protein should be inspected by eye in the first minutes upon dilution and in case clear solution becomes hazy, additional millilitres of dialysis buffer should be added to increase salt concentration.

**Figure 6.**
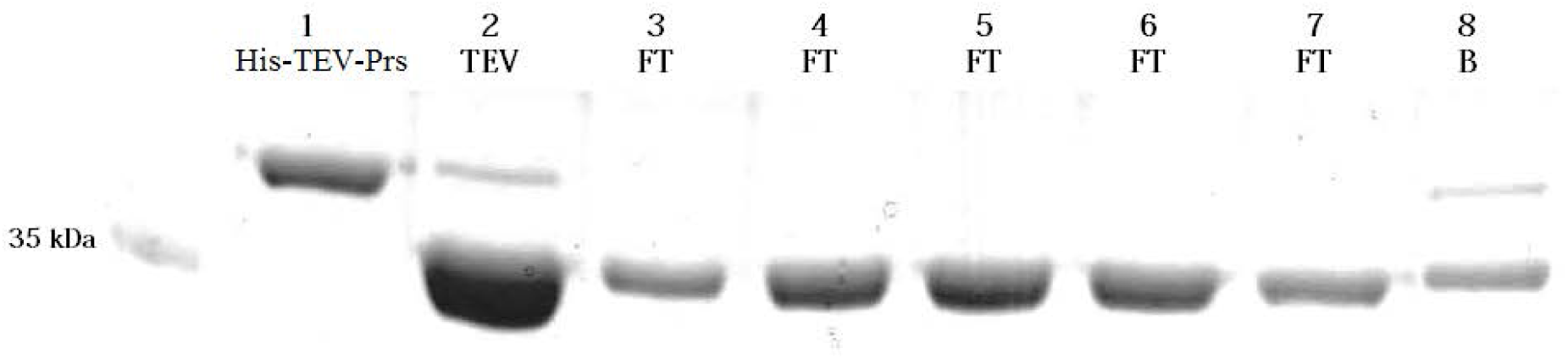
Prs after TEV cleavage and purification. 1: His-TEV-Prs protein; 2: TEV- Prs protein after overnight TEV cleavage in Dialysis buffer, 3-7 FT: untagged Prs eluted in the flow through fraction from the His-Trap column; 8 B: elution with buffer B containing imidazole. The TEV cleavage was approximately 80-85 % efficient. Fractions 2-7 were used for further purification steps. Each fraction was diluted with 2x SDS loading dye and 10 µl was loaded on a 10 % SDS-PAGE gel. After separation, proteins on a gel were visualised with Coomassie Brilliant Blue protein staining.

Untagged Prs after TEV cleavage was successfully concentrated in either SEC1 or SEC3 buffer. On the contrary, SEC2 buffer with lower NaCl salt concentration or HEPES based buffer caused significant protein loss due to protein aggregation on a membrane during concentration. Thus, untagged Prs necessitated high NaCl salt concentration or potassium glutamate to maintain stability.

The protein centrifugation on Millipore Amicon Ultra concentrators is a critical step to sustain high yield of the untagged protein. We found cellulose membrane used in the Millipore concentrators as most efficient, providing the centrifugation temperature is between 16°C and 20 °C for His-Prs constructs and 20 °C for untagged Prs.

Activity of the purified untagged Prs was slightly lower upon dialysis and storage in SEC1 buffer in comparison to untagged Prs in SEC3 buffer and His-Prs (Figure 4A). Thus, although Prs in SEC1 buffer with 500 mM NaCl is relatively stable, we recommend 150 mM L-Glutamic acid monopotassium salt buffering conditions for untagged Prs related assays.

Using the protocol described in this work, we have successfully purified two fusion protein variants of Prs (Figure 7): with short N-terminal His_6_-tag and with cleavable His_6_-tag. We successfully removed the tag from the protein using TEV protease. In addition, we have also purified catalytically inactive His_6_-tagged variant His-PrsK194A and the temperature-sensitive His-PrsG9S protein variant.

**Figure 7.**
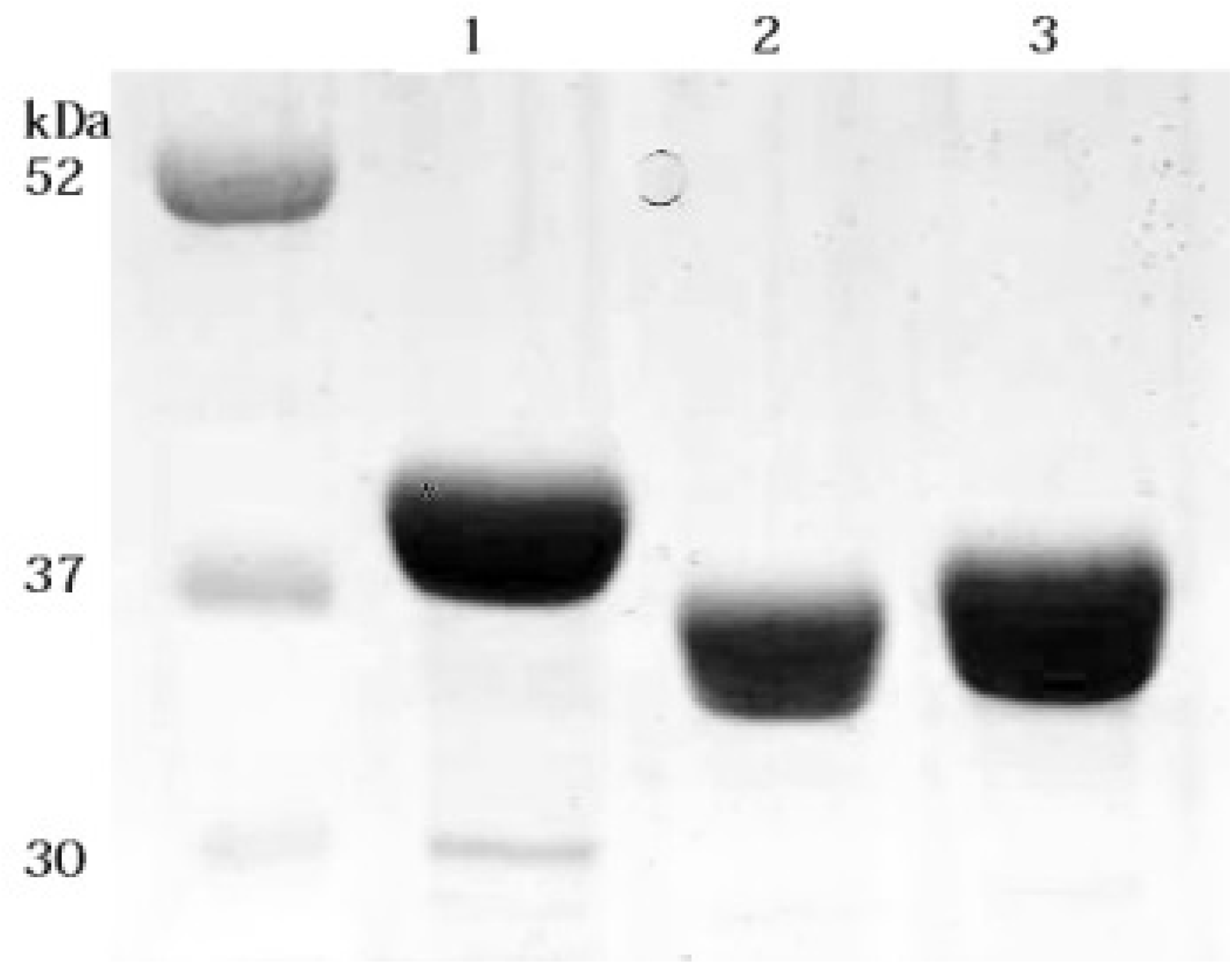
Prs variants purified in this study. 1: His-TEV-Prs; 2: Prs untagged, 3: His-Prs. Calculated protein size was 36.99 kDa, 34.47 kDa, 35.24 kDa respectively. Purified and concentrated protein aliquots were diluted with 4x SDS loading dye and 8 µl was loaded on a 10 % SDS-PAGE gel. The proteins on a gel were visualised with Coomassie Brilliant Blue protein stain solution.

## CONCLUSIONS

We showed here that phosphoribosyl phosphate synthase (Prs) protein from *E. coli* is sensitive to low temperatures during purification process and for highest yield and quality of the protein, purification should be performed at room temperature. We proposed 3 buffer choices providing good stability and activity for His-Prs protein fusion and untagged Prs protein to facilitate buffer selection for further study depending on the conditions desired for downstream applications. Corroborating previously published results, we showed that Prs G9S variant has lower catalytic activity *in-vitro* at permissive temperature. This result confirmed that the *E. coli* PRPP synthase purification protocol described in this work provides active and stable proteins, exhibiting similar properties to those observed in *in-vivo* studies.

## ACKNOWLEDGMENTS

This work was financially supported by the National Science Centre, Poland under grant agreement No. UMO-2014/13/B/NZ2/01139 and UMO-2017/01/X/NZ1/01327 and by the Federation of European Microbiological Societies under FEMS-GO-2017-028 research and training grant.

